# Immunogenic fusion proteins induce neutralizing SARS-CoV-2 antibodies in the serum and milk of sheep

**DOI:** 10.1101/2022.12.11.519990

**Authors:** Gregory M. Jacobson, Kirsty Kraakman, Olivia Wallace, Jolyn Pan, Alex Hennebry, Grant Smolenski, Ray Cursons, Steve Hodgkinson, Adele Williamson, William Kelton

## Abstract

Antigen-specific polyclonal immunoglobulins derived from the serum, colostrum, or milk of immunized ruminant animals have potential as scalable therapeutics for the control of viral diseases such as COVID-19. Enhancing the efficacy of vaccine antigens to induce robust and specific antibody responses remains central to developing highly effective formulations. The direct fusion of immunoglobulin (IgG) Fc domains or other immune-stimulating proteins to antigens has shown promise in several mammalian species but has not yet been tested and optimized in commercially-relevant ruminant species. Here we show that the immunization of sheep with fusions of the receptor binding domain (RBD) of SARS-CoV-2 to ovine IgG2a Fc domains promotes significantly higher levels of antigen-specific antibodies compared to native RBD or full-length spike antigens. This antibody population was shown to contain elevated levels of neutralizing antibodies that suppress binding between the RBD and soluble hACE2 receptors *in vitro*. The parallel evaluation of a second immune-stimulating fusion candidate, Granulocyte-macrophage colony-stimulating factor (GM-CSF), induced high neutralizing responses in select animals but narrowly missed achieving significance at the group level. Furthermore, we demonstrate that the antibodies induced by these fusion antigens are transferred from maternal serum into colostrum/milk. These antibodies also demonstrate cross-neutralizing activity against diverse SARS-CoV-2 variants including delta and omicron. Our findings highlight a new pathway for recombinant antigen design in ruminant animals with applications in immune milk production and animal health.

## Background

The COVID-19 pandemic has focused global efforts to develop new therapies that mitigate the severity of the disease at the individual and population level. While vaccination provides excellent prophylactic protection against the original SARS-CoV-2 virus, the rapid evolution of new variants (e.g delta B.1.617.2, omicron B.1.1.529 and omicron BA.5) and the current inability to vaccinate within certain demographics (e.g. infants) has necessitated the development of alternate strategies to both limit disease transmission and treat COVID patients. One potential solution is to create formulations of neutralizing polyclonal antibodies derived from ruminant animals immunized with SARS-CoV-2 viral proteins. A significant advantage of this approach is the inherent scalability enabled by integration into existing ruminant farming systems.

SARS-CoV-2 viral entry is mediated by spike protein S1 binding to the human angiotensin-converting enzyme receptor (hACE2) through its N-terminal receptor-binding domain (RBD). The distribution of hACE2 receptors on cells defines viral tropism within the host. In humans, these receptors are expressed most abundantly in epithelial cells of the respiratory and digestive tracts (Hamming et al., 2004; Lee et al., 2020). Neutralizing antibodies that block this interaction are demonstrated to provide protection against certain SARS-CoV-2 variants, with FDA approval granted for several monoclonal antibody products (Hwang et al., 2022). While highly effective against early pandemic variants, these monoclonal antibodies have reduced activity to new and emerging SARS-CoV-2 variants as escape mutations arise (Cao et al., 2021), which has led to the FDA revising the authorizations for multiple mAb SARS-CoV-2 approvals (e.g. bamlanivimab and etesevimab, REGEN-COV; casirivimab and imdevimab, and sotrovimab) (Cavazzoni, 2022; Mullard, 2022). This, in turn, has focused attention on alternative treatment strategies with polyclonal or cocktails of monoclonal antibodies as a way of providing broader coverage against SARS-CoV-2 variants of concern (Alape-Girón et al., 2021).

Due to high global demand for COVID-19 treatments, the immunization of ruminant animals has been proposed as a cost-effective approach to produce large quantities of polyclonal SARS-CoV-2 neutralizing antibodies in serum or colostrum/milk (Ainsworth et al., 2020; Arenas et al., 2021; Jawhara, 2020). The resulting hyperimmune fractions could either be formulated directly as a functional food supplement or processed to extract purified pathogen-specific antibodies for direct therapeutic use. Such strategies have previously demonstrated protection against a variety of viral and bacterial pathogens including Middle East Respiratory Syndrome (MERS) (Luke et al., 2016), Human Immunodeficiency Virus (HIV) (Kramski et al., 2012), Rotavirus (Mitra et al., 1995; Sarker et al., 2007), Enterotoxigenic *Escherichia coli* (Savarino et al., 2017) and *Clostridium difficil*e (Steele et al., 2013; van Dissel et al., 2005). In addition to neutralization via steric hindrance of interactions essential for pathogen infection, ruminant antibody fractions may also engage with elements of the human immune system to mediate potent effector functions (Ulfman et al., 2018). Most recently, Kangro et al. immunized pregnant cows with the RBD subdomain of the SARS-CoV-2 spike protein and formulated the resulting colostral antibodies for nasal application (Kangro et al., 2022). Neutralizing antibodies against SARS-CoV-2 were reported to persist in the nasal mucosa for at least four hours in a small clinical trial; a timescale that potentially confers protection against viral infection. This study also demonstrated that the protective capacity of anti-SARS-CoV-2 hyperimmune formulations is closely correlated with the abundance of pathogen-specific neutralizing antibodies induced by immunization. Approaches to improve neutralizing antibody levels in colostrum/milk are of particular importance as titers decline within hours of calving (Elfstrand et al., 2002). Colostral/milk antibodies are accumulated primarily from the systemic circulation, via active FcRn transport, with a more minor contribution from local production in the mammary glands (Hurley & Theil, 2011). Therefore, high antibody titers in colostrum/milk are most likely achieved by strengthening and improving the duration of serum response to antigens.

Several studies have reported the use of immunostimulant technology to improve antigenicity of the SARS-CoV-2 receptor binding domain (RBD) and thereby improve specific antibody titers. The RBD region comprises 222 amino acids, representing around 17% of the full-length viral spike protein (1273 amino acids). Antigens based on this smaller region avoid the expression challenges associated with using heavily glycosylated full-length spike proteins, and may also focus antibody responses towards critical viral neutralizing epitopes (Kleanthous et al., 2021).

Combination or fusion of the RBD with immune-stimulating proteins can further boost antibody levels following immunization. One such fusion partner widely used in non-ruminant species is the IgG Fc domain, which promotes antigen half-life extension and targets the antigen to Fc receptor expressing cells that mediate antigen presentation (Czajkowsky et al., 2012). Multiple murine studies have shown that fusion of the RBD to murine IgG Fc regions induces robust neutralizing responses upon immunization (Liu et al., 2020; Qi et al., 2020). Similar results have also been observed in non-human primates where a human IgG1 Fc RBD fusion vaccine provided protection against challenge with live SARS-CoV-2 virus (Sun et al., 2021). Beyond Fc fusions, the concurrent administration of other soluble immune-stimulating factors including cytokines (e.g. GM-CSF) have also shown promise in enhancing anti-SARS-CoV-2 immunity (Vernet et al., 2021).

Here we report the development of new ruminant-specific antigens for the induction of antibody-based SARS-CoV-2 immunity. In particular, we show the fusion of ovine IgG2a Fc regions to the RBD domain induced high and sustained serum neutralizing titers in sheep. Further, these antibodies persist in ovine colostrum/milk following lambing and have broad SARS-CoV-2 cross-variant neutralizing potential. Our antigen designs have direct application in the production of high potency SARS-CoV-2 hyperimmune milk and could be further adapted as recombinant vaccines for animal health in commercially important ruminant species. To our knowledge this is the first use of immune regulatory elements incorporated in to antigenic fusion proteins to improve the nature of the immune response in ruminants.

## Methods

### Antigen production

#### Antigen design

Antigen designs are summarized in the main text (Fig. 1a, b), with sequence data provided as a supplement (Additional file 1: Tables S1 and S2). 14,429 spike protein sequences were downloaded from the GSAID database (Shu & McCauley, 2017) (accessed: 10 Aug 2020) and aligned to obtain a consensus sequence. All antigen designs were based on this consensus sequence, which mapped to the original Wuhan SARS-CoV-2 sequence with an additional D614G mutation. Ovine IgG2 Fc cDNA sequences (Clarkson et al., 1993) were obtained from the IMGT database (Lefranc, 2011) (IMGT accession number: X70983, bases 313-987). The nucleotide sequence of ovine GM-CSF was back-translated from the amino acid sequence obtained from the UniProt database (UniProt accession number: P28773, amino acids 18-144). Each design was appended with an N-terminal rabbit IgH signal peptide sequence for soluble expression and a C-terminal 6His tag for downstream purification.

**Figure 1.**
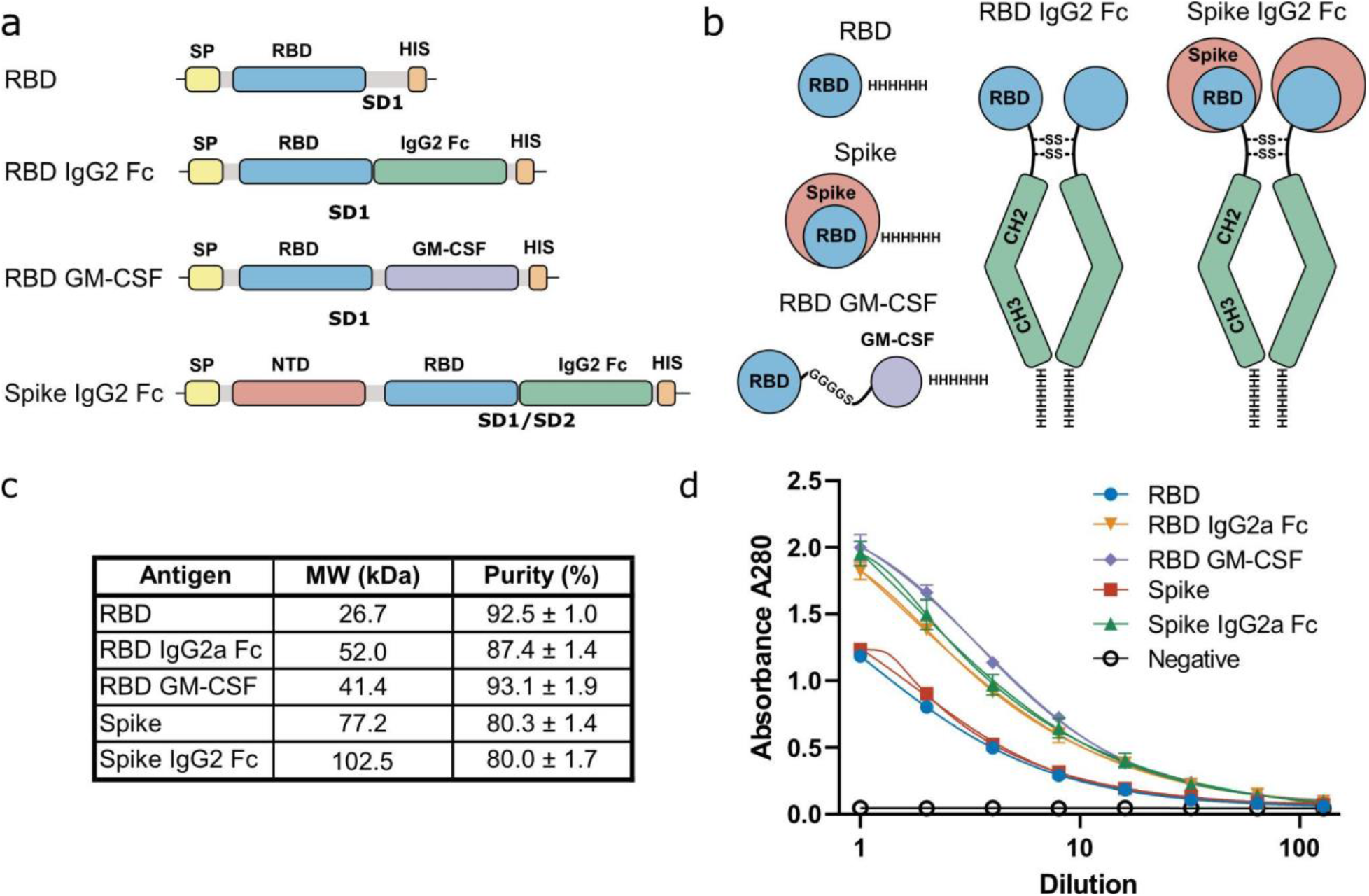
Design and characterization of ovine-specific SARS-CoV-2 antigen fusions. **a** Schematic of designs at the DNA level showing C-terminal immunostimulatory domains IgG2a Fc or GM-CSF. **b** Schematic of designs at the protein level showing dimerization of the IgG2a Fc fusion proteins. **c** Summary table showing antigen purity as determined by SDS-PAGE gel followed by densitometry. Purities are listed with standard error calculated from three replicates loaded at 1, 2, and 4 μg of antigen. **d** ELISA analysis of antigen binding to soluble hACE2 HRP receptors with Ovalbumin used as a negative control.

#### Protein expression

Antigen plasmid constructs were synthesized commercially and provided in the pTwist-CMV-BetaGlobin-WPRE-Neo vector (Twist Biosciences). Heavy and light chain plasmid sequences for producing monoclonal antibodies LY-CoV555, REGN-33, and REGN-87, were a kind gift from Professor Sai Reddy (ETH Zurich). DH5α *E. coli* stocks for each construct were produced by transformation into chemically competent cells. Transfection-ready plasmids were purified from bacterial cultures by maxiprep (Invitrogen, #K20006). Expi293 HEK cells (Thermofisher, #A14527) were expanded and passaged in Expi293 expression media (Thermofisher, #A1435101) according to the manufacturer’s recommendations. Briefly, cells were grown in suspension at 125 RPM on a shaking platform (Thermofisher, #88881102) at >80% humidity, 37 °C and 8% CO_2_. Cells were transfected at a density of 3×10^6^ cells/mL with 1 μg of plasmid per mL of culture according to the manufacturer’s instructions. Monoclonal antibody plasmids were mixed at a 2:1 light chain to heavy chain ratio prior to transfection. Supernatants were harvested at day 6 post-transfection by centrifugation of the culture media for 20 min at 6000 g. The cell pellet was then discarded, and the centrifugation step repeated to further remove cell debris. Each supernatant was then filtered through 0.22 μm filters (Corning, #430773) in preparation for purification.

#### Protein purification

Antigen purifications were performed on a BioRad NGC chromatography system. Endotoxin was removed from the system by soaking overnight in 1 M sodium hydroxide. With the exception of omicron B.1.1.529 RBD, which was obtained commercially (Acro Biosystems, USA), all RBD variants (wuhan, beta B.1.351, delta B.1.617.2), and RBD GM-CSF antigens were purified by Ni-NTA affinity chromatography. Supernatants from each culture were dialyzed into 1.5 L 1X PBS for 24 hours using 6-8 kDa molecular weight cut off (MWCO) dialysis tubing (Repligen, #132655). Dialysed samples were collected and imidazole was added to a final concentration of 20 mM. Each sample was then loaded onto a 5 mL HisTrap FF Ni-NTA column (Cytiva, #17525501) pre-equilibrated with 10 column volumes (CV) 1X PBS containing 20 mM imidazole (Buffer A). The column was washed with 5 CV Buffer A before elution with a 4-100 % gradient of 1X PBS containing 500 mM imidazole over 5 CV. Spike protein was provided by Dr Campbell Sheen (Callaghan Innovation, New Zealand).

Ovine IgG2 Fc fusions (RBD IgG2 Fc, Spike IgG2 Fc) were purified by Protein G affinity chromatography using 5 mL HiTrap Protein G HP columns (Cytiva, #17040501). Supernatants were directly loaded onto columns pre-equilibrated with 10 CV 50 mM Na_2_PO_4_, pH 7.4 (Buffer C). After 5 CV of washing with Buffer C, sample elution was performed using a 0-100% 0.1 M formic acid, pH 2.5 gradient over 5 CV. Following elution, fraction purity was assessed by SDS-PAGE gel electrophoresis (BioRad, 12% Criterion XT #3450118, BioRad NZ Ltd). Pooled fractions were buffer exchanged into 1X PBS using 10 kDa MWCO centrifugal spin concentrators (Merck, Cat#UFC801024), for 15 min at 4000 g. Antigens were adjusted to ∼0.5 mg/mL with sterile 1X PBS (pH 7.4) and stored at -80 °C.

Monoclonal antibody controls were purified via gravity flow using Protein G Sepharose resin (Life Technologies, #101243). Filtered supernatants were passed three times through gravity flow columns pre-equilibrated in 1X PBS. After 5 CV of column washing, antibodies were eluted with 4 mL 100 mM formic acid pH 2.0 and immediately neutralized with 500 μL of NH_4_OH solution. Antibodies were buffer exchanged back into 1X PBS using 10 kDa MWCO centrifugal spin concentrators, normalized to 1 mg/mL and stored at -20 °C.

### Antigen characterization

#### Antigen purity

Antigen samples of 1, 2, and 4 μg were run on pre-cast gels (12 % Criterion XT #3450118, BioRad NZ Ltd). Gels were stained in Fairbanks A and de-stained in 10 % acetic acid before image capture using a Bio-Rad GS900 Calibrated Densitometer. Image Lab software (BioRad) was used for automatic band detection. Sample purity was determined as percentage of the target protein band density to the total protein density in the lane. The total protein purity was calculated as an average of 1, 2, and 4 μg sample loadings for each antigen.

#### ACE2 Receptor Binding

Antigen functional activity was confirmed by assessing ACE2 binding relative to a spike protein positive control (Callaghan Innovation, New Zealand) and ovalbumin negative control protein (Sigma, #A5503). ELISA plates were coated for 1 hour by incubation at ambient temperature with 50 μL antigen (0.072 nM) in 50 mM Na_2_CO_3_, pH 9.5. Plates were blocked for 1 hour at room temperature with 1X PBS with 2% skim milk powder and 0.05% Tween (PBSMT). Plates were then washed six times with 1X PBS with 0.05% Tween (PBST) before 120 μL of ACE2-HRP protein (ProteoGenix, France) at a 1:200 dilution in 1X PBS containing 2% milk powder (PBSM) was added to the first well of each plate column, and serially diluted 1:2 down the plate into wells containing 60 μL PBSM. Plates were incubated at ambient temperature for 1 hour and washed six times with PBST before the addition of 50 μL TMB solution. After 10 min, the reaction was quenched with 50 μL 1 M H_2_SO_4_ per well and the plates read at 450 nm (and 570 nm for background). All ELISA curves were fitted with a five-parameter asymmetric logistic model (Richards equation) in Graphpad Prism 9.3.0.

### Animal Trial

#### Animal cohort

Regulatory approvals were obtained from the Ruakura Animal Ethics Committee (approval #15191) and MPI ACVM (agricultural compounds and veterinary medicines). An initial cohort of 100 non-pregnant hoggets (∼12 months in age) at the Ruakura Research Campus (New Zealand) was evaluated for use in the trial. Animals with weights of ∼52 kg were selected, with flighty animals, and those with noticeable health issues such as coughing or lameness, excluded. The animals were randomly allocated to eight trial groups each comprising seven animals. The sheep were maintained on normal pasture rations throughout the trial, with water and pasture available *ad libitum*. For day-to-day care, the sheep were managed by AgResearch Ltd (New Zealand) staff.

#### Immunogen preparation

Immunogens were prepared as water in oil emulsions on the day of injection. Briefly, Freund’s Incomplete Adjuvant (Sigma, # F5506-10 × 10 mL; Lot #SLCD0710) and sterile PBS were aliquoted into sterile 50 mL tubes the day prior to immunization and stored at 4 °C overnight. Antigens were also taken out of the -80 °C freezer and allowed to thaw overnight at 4 °C. A total of 18.45 mL of Freund’s Incomplete Adjuvant was combined at a 4:1 ratio with each trial antigen (in PBS up to 6.15 mL) and 0.123 mL 1% v/v Triton X-100 detergent in a sterile 50 mL tube to make a total volume of 24.6 mL of each immunogen mixture. Antigens were concentration corrected to give molar amounts of antigen equivalent to ∼3.3 nmol with the exception of the Spike IgG2 Fc antigen which was normalized to ∼1.3 nmol. A second “Spike low” cohort was included at the same 1.3 nmol concentration for comparison. Emulsification of immunogen was achieved using an Ultra Turrax Homogeniser fitted with an 8 mm probe for 5 minutes in a 50 mL polypropylene tube. Emulsification was conducted on ice with frequent mixing to prevent local heating of the mixture. Emulsions were then stored at 4 °C before use. The molecular equivalent masses for antigens are given as a supplement (Additional file 1: Table S3).

#### Trial sampling

In the first phase of the trial, serum responses to each of the SARS-CoV-2 antigens were evaluated. Hoggets (12 mo, median weight 52 kg) were immunized three times on days 0, 21, and 42 over the 16 week trial period with the study antigens (Fig. 2A). Antigen doses were injected (1.4 mL) at shaved sites on the hind leg. Blood samples were collected in BD vacutainers (BD, #367895) on days 0, 9, 21, 42, 63, 84, and 112. Each sample was centrifuged at 1200 g for 15 min at 4 °C and the serum fraction collected for storage at -20 °C. The trial was subsequently continued in a second phase with a restricted number of cohorts to evaluate the transfer of anti-SARS-CoV-2 antibodies to colostrum/milk (Fig. 4A). Ewes were mated and pregnancy was confirmed before a fourth immunization ∼3 weeks before lambing. Immediately after lambing, both colostrum and blood samples were collected with subsequent samples taken on days 4, 16, and 32. All samples from both trial phases were aliquoted and stored at -20 °C prior to analysis of anti-SARS-CoV-2 activity.

**Figure 2.**
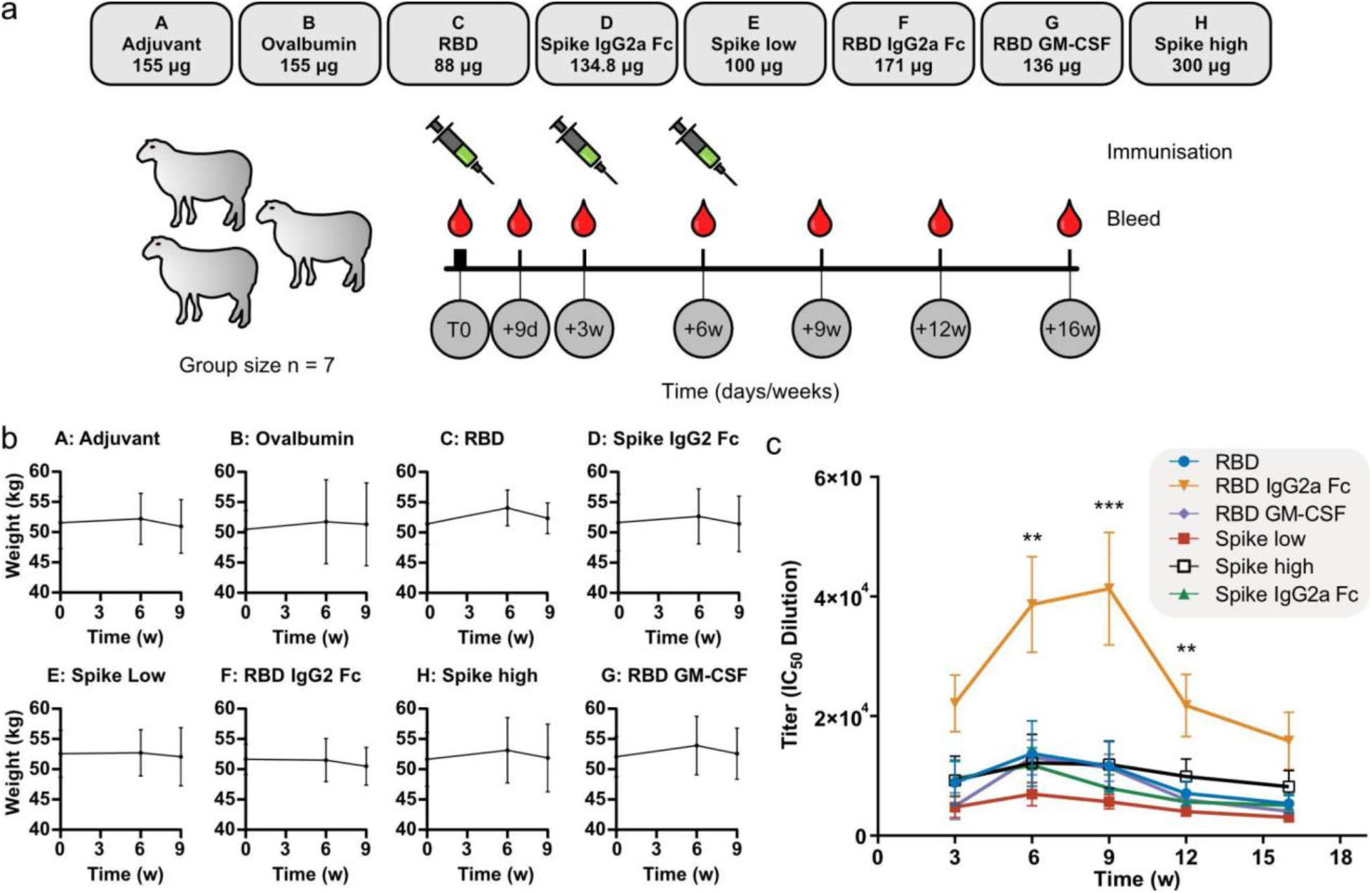
Evaluation of anti-SARS-CoV-2 antibody responses in ovine serum following immunization with designed antigens. **a** Schematic showing the design of a 16 week ovine trial to test designed antigen immunogenicity. Hoggets were randomly allocated to groups (n = 7), immunized once and boosted twice with SARS-CoV-2 antigens over a six week period. Adjuvant-only and Ovalbumin groups were included as negative controls. Blood was periodically sampled for serum analysis until 16 weeks from the first immunization. **b** Animal weights in each group during the immunization phase of the trial. **c** ELISA analysis of anti-SARS-CoV-2 RBD (Wuhan) titers for each group during the trial. No detectable response was observed for either Adjuvant or Ovalbumin control groups. Error bars show the standard error within each group. ANOVA testing was used to measure the statistical significance between groups. * p < 0.05, ** p < 0.01, *** p < 0.005 are shown only for the RBD IgG2a Fc group relative to the RBD group.

#### Serum titer ELISA

High-binding ELISA plates (Corning, #3590) were coated by overnight incubation at 4 °C with 100 μL prepared RBD-His antigen (1 μg/mL in 0.05 M carbonate bicarbonate buffer, pH 9.6). Following the binding incubation period, plates were washed three times with 300 μL wash solution (0.05 M Tris base, 0.14 M NaCl, 0.05% Tween20, pH 7.5; TBS-T) and blocked by shaking incubation with 200 μL blocking solution (5% horse serum in TBS-T) for 1 hour at ambient temperature. Blocked plates were washed three times with the wash solution. Serum samples for analysis were diluted in the blocking solution to provide six dilution points per sample. For some samples, only five points were available due to matrix interference at low dilutions. A reference curve was also included on each ELISA plate along with a blank. All samples were assessed in duplicate. Diluted serum samples (100 μL) were added to wells and incubated with shaking for two hours at ambient temperature. Incubated plates were washed five times with 300 μL of wash solution. HRP-conjugated mouse anti-sheep IgG secondary antibody (Jackson #213-032-177) (100 μL) made up at 1/5000 dilution in blocking solution was added to each well. Plates were wrapped in foil and incubated with shaking at ambient temperature for one hour. Incubated plates were washed five times with 300 μL of wash solution. TMB reagent (Surmodics, #TMBW-1000-01) reagent (100 μL) was added to each well prior to shaking incubation at ambient temperature with the plate covered in foil. After 15 min, the reaction was quenched by the addition of 50 μL 2 M H_2_SO_4_ to each well, and the plate was read at 450 nm and 600 nm with a five-second shaking step before analysis. As with ACE2-HRP ELISA curves, IC50 values were determined by fitting a five-parameter logistic model in Graphpad 9.3.0 and reported with standard errors. Statistical differences between groups were evaluated by ANOVA with correction for multiple comparisons.

#### Surrogate viral neutralization assay (sVNT)

ELISA strip wells (Corning, #2580) were coated by overnight incubation at 4 °C with 75 μL native, beta, delta, or omicron RBD-His antigens at 2.67 μg/mL in 0.05 M NaHCO_3_, pH 9.5. Plates were blocked for two hours at ambient temperature (or overnight at 4 °C) with PBSMT. Blocked plates were washed three times with 360 μL of PBST. Serum or colostrum/milk samples were prepared at 1:5 dilutions in PBSM. All samples were assessed in duplicate. An 8-point serial dilution of 1:3 was performed in PBSM to generate sample volumes of 60 μL. 60 μL ACE2-HRP reagent prepared at 1:750 dilution in PBSM was added to all dilutions and mixed thoroughly. 100 μL of this mixture was added to the plate wells and incubated at ambient temperature with shaking for 1 hour. Plates were washed three times with 360 μL of PBST and 50 μL TMB solution was added for 5 min during which time the plates were incubated in the dark. The reaction was quenched by the addition of 50 μL 1M H_2_SO_4_ to each well, and the plate was read at 450 nm and 600 nm. 450 nm Absorbance values were corrected using the 600 nm (background) reading and the percentage inhibition was calculated as:*% Inhibition* = (*1* − *A450 Sample* / *A450 Negative Control*) × *100*. The resulting curves were fitted to a five-parameter logistic model using Graphpad 9.3.0. Comparisons between treatment and control groups were assessed by ANOVA with correction for multiple comparisons.

#### Mass spectrometry antibody subclass profiling

Colostrum samples were thoroughly mixed, and 2 mL samples were centrifuged at 17,000 g for 10 mins at 4 °C to separate the protein-rich supernatant from the unwanted fat component. Ovine IgG1, IgA and IgM concentrations were determined by LC-MS/MS using a stable isotope dilution assay with heavily-labelled internal peptide standards (Chambers et al., 2022). Briefly, a known volume of colostrum sample was digested overnight at 37 °C with trypsin and internal standards. Target peptides specific to each immunoglobulin isotype were separated by reverse-phase HPLC on a Hypersil Gold C18 column (100mm × 2.1mm; 1.9 μm) and detected using a TSQ Altis triple-quadrupole mass spectrometer (ThermoFisher). The signal intensity of the product ions from the targeted peptides was compared to the corresponding signal intensity of synthetic tryptic peptides of known abundance from a 9-point calibration curve.

## Results

### Design of and production of ruminant specific SARS-CoV-2 antigens

We designed a series of His-tagged antigens that linked ovine IgG2a Fc domains or ovine GM-CSF cytokine sequences to the C-terminus of SARS-CoV-2 RBD or full-length spike protein (Figure 1a, b; Supplementary Table S1, Additional File 1). Due to incomplete information about functional ovine IgG2a hinge region sequences, we performed an alignment to the human IgG1 sequence and selected ovine residues that align to the “DKTH” motif (residues 221 -224 by the Eu numbering system), which we have previously used to generate functional Fc domains that retain Fc receptor binding (Jung et al., 2012; Kelton et al., 2014). The unstructured hinge region of IgG2a Fc was hypothesized to act as a flexible linker sequence between the antigen and the Fc domain. We further modified the full-length spike antigen sequence to remove the N-terminal furin site to prevent cleavage of the Fc fusion upon expression. In parallel, we designed an RBD fusion with ovine GM-CSF by incorporating the full ovine GM-CSF coding region except for the N-terminal signal sequence (residues 18-144). The construct included a GGGGS linker sequence between the RBD domain and the GM-CSF to prevent steric hindrance influencing antigen or cytokine function (Chen et al., 2013). To mediate soluble expression in mammalian cells, we added a rabbit Immunoglobulin Heavy Chain signal peptide sequence at the N-terminus of each design.

Antigen purity exceeded 80% for all antigens when analysed by SDS-PAGE gel and scanning densitometry (Figure 1c; Supplementary Figure S1, Additional File 1). We used Protein G resin as a conformational ligand for purification of IgG2a Fc fusions that additionally confirmed the correct dimerization of these domains. All other antigens were purified by Ni-NTA affinity chromatography. As an additional verification of functional antigen production, we evaluated binding to recombinant hACE2 receptors by ELISA. All purified antigens showed considerably higher binding to hACE2 compared to Ovalbumin control protein (Figure 1d). We also observed a stronger signal from the antigen fusions (RBD IgG2 Fc, Spike IgG2 Fc, RBD GM-CSF) than from the unfused antigens (RBD, Spike), presumably due to a greater availability of hACE2 binding epitopes in native conformations following hydrophobic plate coating.

### RBD-Fc fusions induce elevated levels of neutralizing anti-SARS-CoV-2 antibodies in ovine serum

The ability of our antigen designs to induce high anti-SARS-CoV-2 titers in the serum of ruminant animals was investigated using sheep as a model ruminant species. Here, we randomly allocated groups of hoggets (median age 12 months; median weight 52 kg; n= 7) into eight groups for immunization with the designed antigens (RBD IgG2 Fc, Spike IgG2 Fc, RBD GM-CSF), native antigens (RBD, Spike high, Spike low) and controls (Ovalbumin, and adjuvant alone) (Figure 2a). Antigens were normalized by both molecular weight and purity such that 0.3 nmol of total antigen was injected into each animal, with the exception of group E where the yield of Spike IgG2a Fc allowed only 0.1 nmol of antigen per injection. For this reason, an additional control group of spike protein alone (Group E, Spike low) was included as a control for this group. Our normalization scheme resulted in each group receiving different masses of antigen but an equivalent number of neutralizing SARS-CoV-2 epitopes. Animal health was tracked by physical examination during each blood draw and via animal weights over time (Fig. 2b). Immunization was well-tolerated; a slight, but non-significant, decrease in animal weight was observed at week nine, though this was consistent across all groups including those dosed with control antigens.

To quantify the magnitude of the serum response, we analyzed polyclonal antibody responses against RBD antigen by ELISA screening (Figure 2c; Supplementary Figure S2, Additional File 1). These assays revealed a significant 3.1-fold increase in anti-RBD titer for the RBD IgG2a Fc antigen design compared with RBD antigen alone, with peak antibody responses observed three weeks after the second boost at week nine (IgG2 Fc titer IC_50_ = 4.1 × 10^4^ ± 0.9 × 10^4^, RBD titer IC_50_ = 1.2 × 10^4^ ± 0.4 × 10^4^, p = 0.0038) (Supplementary Figure S3, Additional File 1). Antibody responses persisted until the 16-week time point with titers ∼40% of the peak response (RBD IgG2a Fc titer IC_50_ = 1.6 × 10^4^ ± 0.5 × 10^4^). By comparison, no significant difference in titer was observed between Spike IgG2 Fc and Spike antigen alone, although this is likely due to the measurement of anti-RBD responses rather than anti-spike responses by ELISA. Likewise, the RBD GM-CSF antigen did not elicit antibody titers significantly above the RBD control suggesting lesser immunostimulation by this fusion. No RBD binding signal was observed for either the adjuvant-only or the ovalbumin control groups.

As RBD titers do not provide a direct estimate of neutralization potential, we performed surrogate viral neutralization assays with a dilution series of the hyperimmune sera (Figure 3a; Supplementary Figure S4, Additional File 1). The clinical neutralizing antibody LY-CoV555 antibody was included as a positive control. Compared to the variation in our positive control samples, a large degree of variation is observed predominantly within the RBD IgG2a Fc and RBD GM-CSF cohorts, with multiple high-responding animals mixed with those considered as low-responders (neutralization titers below the positive control signal). In contrast to RBD antibody titers in serum (Figure 2c), we found the highest neutralization capacity in multiple cohorts was generated at week 6 rather than week 9 (Figure 3b; Supplementary Figure S3, Additional File 1). We also observed that the RBD IgG2a Fc immunized cohort had consistently higher neutralization titers over a range of time points compared to the RBD antigen alone cohort, with IC_50_ values of 282 ± 79 verses 83 ± 28 (a 3.4-fold increase; p = 0.02) at week six and 168 ± 55 verses 52 ± 18 (a 3.2 fold increase; p = 0.03) at week nine.

**Figure 3.**
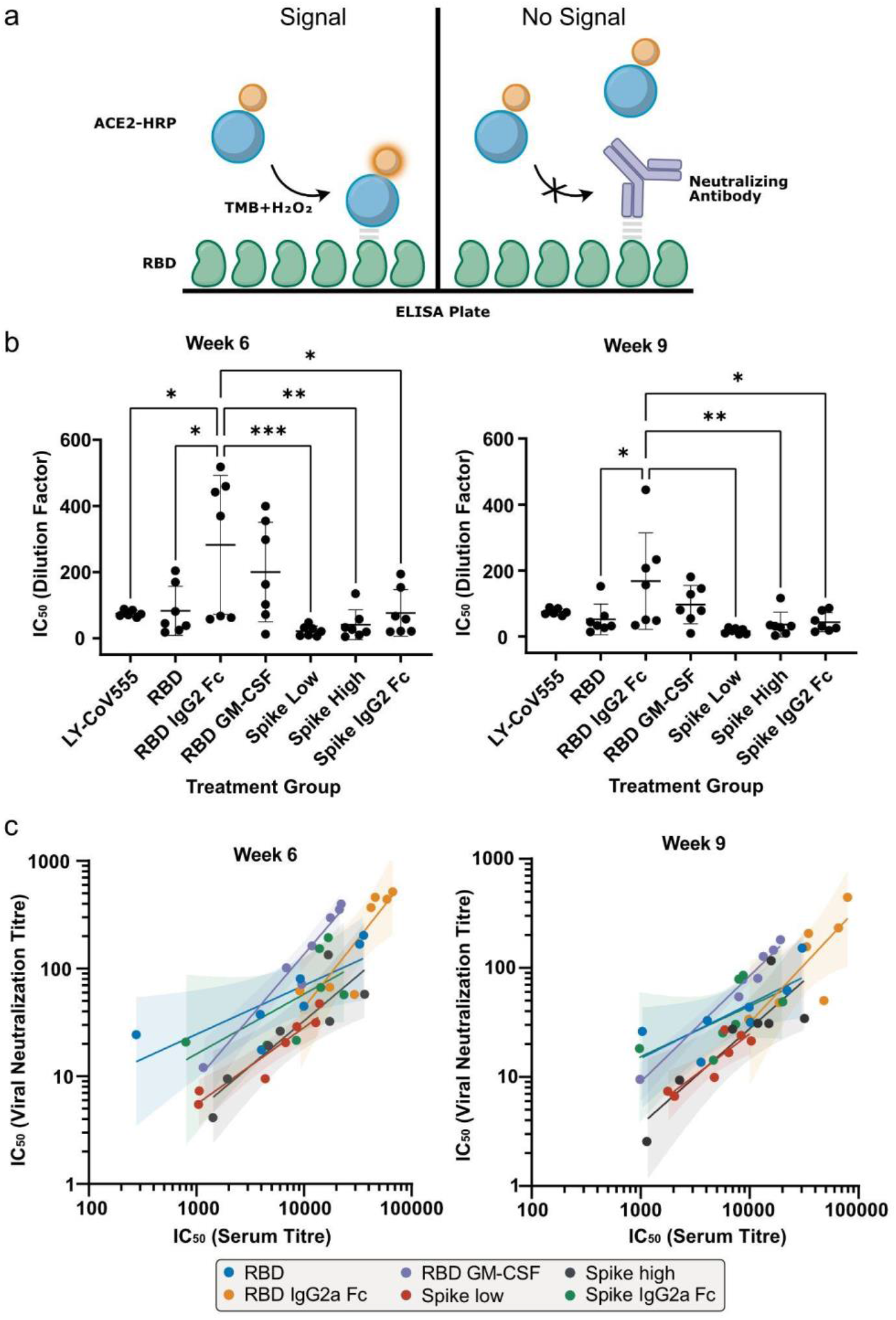
Evaluation of SARS-CoV-2 neutralizing responses in serum by surrogate viral neutralization testing. **a** Cartoon showing the principle of the surrogate viral neutralization test. Neutralization generates a loss of ELISA signal as hACE2-HRP binding to SARS-CoV-2 RBD is reduced by antibody blocking, with such approaches comparable in specificity and sensitivity to live viral assays (Bayarri-Olmos et al., 2021; Tan et al., 2020). **b** Plots showing relative inhibitory concentrations for RBD interaction with hACE2 for each trial group at weeks 6 and 9 as measured by serum dilution factor. LY-CoV555 is the positive control anti-SARS-CoV2 clinical antibody bamlanivimab diluted from 6.9 μg/mL. Error bars show the standard error of each group. ANOVA testing was used to measure the statistical significance between groups with * p < 0.05, ** p < 0.01, and *** p < 0.005. **c** Correlation plot of log transformed anti-SARS-CoV-2 neutralization titer data from the sVNT vs serum titer data against SARS-CoV-2 RBD for weeks 6 and 9. Shaded areas indicate the 95% confidence intervals for power law fitting.

To further investigate whether individual antigens polarize the antibody response towards neutralizing epitopes within the RBD, we first performed Pearson correlation testing on log-transformed serum titer and serum neutralizing titer data for each cohort (Figure 3c). The regression generated power law coefficients that generally centred around 1 (Supplementary Table S2, Additional File 1), indicating proportional increases in neutralizing antibody levels as anti-SARS-CoV-2 antibody titers increased. We then tested to see whether mean neutralizing titers, when corrected for serum titers, were significantly higher for any particular antigen group (Supplementary Table S3, Additional File 1). While some individual animals in the RBD GM-CSF group demonstrated high levels of neutralizing antibodies (D718, D727, D755), there were no significant differences between the group means at either week 6 or 9 (Week 6 RBD GM-CSF vs RBD p = 0.11, Week 9 RBD vs GM-CSF p = 0.099).

### Anti-SARS-CoV-2 antibodies are trafficked into colostrum and milk

While polyclonal antibodies can be extracted from ruminant serum, colostrum and milk are readily obtained non-invasively at scale. Based on the highest observed anti-SARS-CoV-2 neutralizing titers in serum, we selected the RBD IgG2a and RBD GM-CSF cohorts for analysis of anti-SARS-CoV-2 antibody transfer into colostrum/milk. RBD and adjuvant-only groups were selected as positive and negative controls respectively. Single low-responding animals from the serum trial were excluded from each of the RBD GM-CSF and RBD cohorts. Sheep were mated and then boosted with antigen approximately three weeks before lambing (Figure 4a). Of the 26 animals carried through into the second phase of the trial, two further animals from the RBD GM-CSF (D716) and RBD IgG2a cohorts (D743) died or were euthanized for health reasons unrelated to the trial. Periodic blood and colostrum/milk samples were collected up to 32 days after lambing. After boosting, serum antibody titers against the RBD antigen were sustained at similarly high levels to those observed in the initial phase of the trial (RBD IgG2a Fc 3.1 × 10^4^ ± 1.7 × 10^4^ at day 251) (Figure 4b, Supplementary Figure S5, Additional File 1).

**Figure 4.**
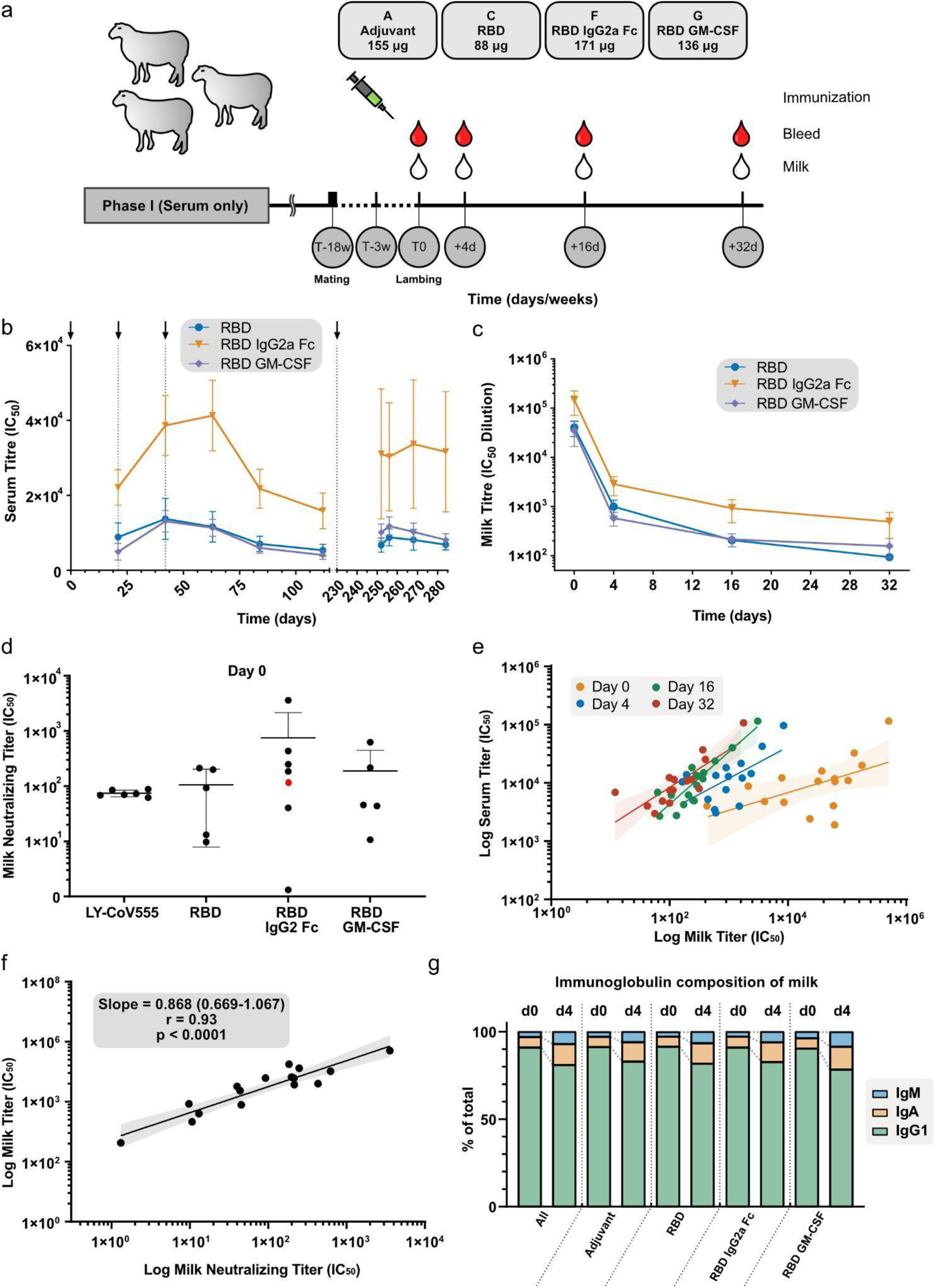
Analysis of anti-SARS-CoV-2 antibodies trafficked into colostrum and milk. **a** Schematic showing the continuation of the ovine trial with select groups. Sheep were mated before a single immunization booster with blood and colostrum/milk samples collected immediately postpartum followed by periodic sampling over 32 days. **b** ELISA analysis of anti-SARS-CoV-2 RBD (Wuhan) titers in serum for each group over both phases of the ovine trial with black arrows indicating immunization timepoints. **c** ELISA analysis of anti-SARS-CoV-2 RBD titers in first colostrum and milk. **d** Neutralizing antibody titers in first colostrum by sVNT at day 0 postpartum. The point in red was not included in the analysis due to the animal being euthanized in the trial. **e, f** Correlation plots of log transformed anti-SARS-CoV-2 RBD serum titer data against anti-SARS-CoV-2 RBD colostrum/milk data (**e**) and log transformed anti-SARS-CoV-2 RBD colostrum/milk titer data against anti-SARS-CoV-2 colostrum/milk neutralization titer data from the sVNT (**f**). The r value indicates the Pearson correlation coefficient. **g** Overview of the change in immunoglobulin subclass composition in colostrum/milk between days 0 and 4 postpartum as measured by mass spectrophotometry. Error bars for all plots show the standard error of each group.

Anti-RBD antibody titers were highest in RBD IgG2a Fc immunized animals at 1.5 × 10^5^ ± 0.8 × 10^5^ in the first colostrum and rapidly decreased by day 4 in all cohorts (Figure 4c, Supplementary Figure S5, Additional File 1). The decline in titer coincided with a sharp drop in total antibody concentration from 75.6 ±12.0 mg/mL at day 0 (colostrum) to 4.8 ± 0.7 mg/mL at day 4 (milk) as determined by mass spectrometry analysis and was expected with the transition from the colostrum to milk phases of the lactation (See Supplementary Figure S6, Additional File 1). We then used surrogate viral neutralization assays to evaluate the neutralizing capacity of colostrum taken at day 0 (Figure 4d, Supplementary Figure S7, Additional File 1). Colostrum samples from individual animals were diluted fivefold and compared against the LY-CoV555 positive control antibody. A large variation in individual animal responses resulted in non-significant differences between the cohorts despite several very high responding animals observed in the RBD IgG2a Fc cohort (IC_50_ 749 ± 571 RBD IgG2a Fc vs 106 ± 44 RBD alone; p = 0.5). To establish broader trends as to how serum titer translates into antibody transfer to the colostrum/milk we fitted power law functions to log transformed titer data (Figure 4e, Supplementary Table S4, Additional File 1). For each day analyzed (0, 4, 16, 32), significant correlations (p≤0.05) were observed that indicate greater antibody accumulation in the colostrum/milk is a function of higher serum antibody concentration. However, we observe steeper gradients at later sampling timepoints (Day 16, 32) suggesting an accumulation of anti-SARS-CoV2 antibodies in early colostrum is not as sustained as milk production begins. The correlation between colostrum anti-RBD titer and colostrum neutralizing potential at Day 0 was also highly correlated with an r value of 0.93; p<0.0001 (Figure 4f).

We finally sought to establish whether the fusion of immunomodulatory domains to RBD antigens alters the immunoglobulin subclass profile in colostrum or milk. IgM, IgA and IgG1 levels (including both antigen specific and non-specific populations) were quantified by mass spectrometry (Figure 4g). In colostrum averaged across all cohorts on day 0, 91.0 ± 0.5% of the total immunoglobulins were IgG1, which reduced after the first four days to 81.4 ± 1.2%. The observed IgM and IgA proportions increased from 3.0 ± 0.2% and 5.9 ± 0.4% to 6.7 ± 0.6% and 11.8 ± 1.0% respectively (Figure 4g). No significant differences in antibody subclass ratios were observed between the cohorts indicating that the antigens did not bias the antibody subclass composition in colostrum/milk.

### Hyperimmune colostrum has broad cross-variant neutralizing potential

It is now well established that human vaccines containing native (Wuhan) spike protein sequences are less effective at generating protective immunity against emerging SARS-CoV-2 variants of concern (Tseng et al., 2022). We therefore evaluated the ability of pooled colostrum (Excluding low-responding animal D708) from the RBD IgG2 Fc cohort to cross neutralize native, beta, delta, and omicron RBD variants using our established surrogate viral neutralization assays. Clinical antibodies REGN-33 and REGN-87 were included as controls that bind distinct RBD epitopes (Figure 5a). REGN-33 performed comparably well against the native and delta RBD with average IC50s of 0.41 ± 0.03 μg/mL and 0.42 ± 0.03 μg/mL respectively but had significantly reduced activity against the beta (average IC50 1.5 ± 0.22 μg/mL) and omicron (average IC50 not determined) RBDs (Figure 5b). In contrast, REGN-87 performed comparably against all the RBDs tested except omicron where IC50 value were not reliably determined (average IC50s; native 0.53 ± 0.12 μg/mL, beta 0.91 ± 0.08 μg/mL, delta 0.40 ± 0.06 μg/mL). Hyperimmune colostrum provided broader cross-neutralization than any individual monoclonal antibody although required high antibody concentrations in the mg/mL range presumably due to only a fraction of the total immunoglobulins being RBD specific (average IC50s; native 0.27 ± 0.01 mg/mL, beta 0.50 ± 0.01 mg/mL, delta 0.21 ± 0.01 mg/mL, omicron 0.48 ± 0.03mg/mL). We additionally observed low levels of naive colostrum neutralizing activity against native, delta and omicron RBD domains despite this cohort receiving only adjuvant during immunization.

**Figure 5.**
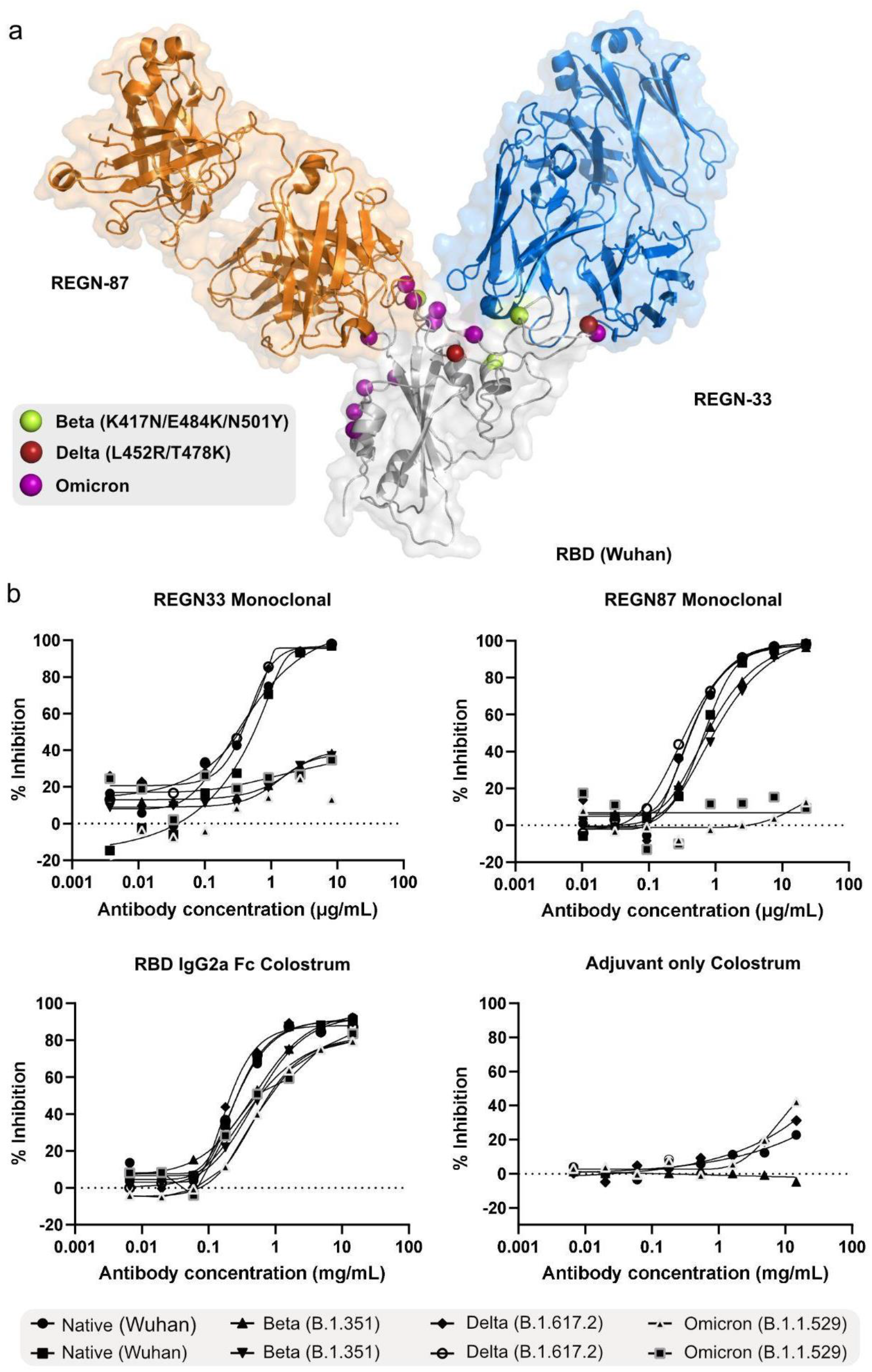
SARS-CoV-2 cross variant neutralizing capability of colostrum pooled from sheep immunized with RBD IgG2a Fc. **a** PBD image 6XDG (Hansen et al., 2020) showing the association of clinical neutralizing antibodies REGN-33 (casirivimab) and REGN-87 (imdevimab) with SARS-CoV-2 RBD. Point mutations in the RBDs of beta and delta SARS-CoV-2 variants are indicated. **b** Relative inhibition of SARS-CoV-2 variants by REGN-33, REGN-87, or pooled RBD IgG2a Fc group colostrum as measured by sVNT. Total colostrum antibody concentrations were determined by mass spectrometry analysis. Percent inhibition was calculated by normalizing each curve to the adjuvant only negative control colostrum.

## Discussion

The production of polyclonal antibodies in ruminant animals has long held promise as a scalable source of cost-effective therapeutics and diagnostic reagents. Maximizing the production of pathogen-specific neutralizing antibodies is critical to producing high-efficacy formulations derived from serum or colostrum/milk. In this study, we increased antibody titers in sheep by enhancing antigen immunogenicity through the fusion of IgG2a Fc domains derived from the host immune system. Using SARS-CoV-2 RBD subunit model antigens, we observed elevated levels of serum antibodies that were subsequently transported into milk following lambing.

Polyclonal antibody preparations from sheep (Dart et al., 1997), and horses (Boyer et al., 2009) remain the standard of care for envenomation and are approved clinically as single-use heterologous injectables with neutralizing activity. In most cases, F(ab’)_2_ molecules are generated to reduce unwanted immunogenicity and prevent other unwanted inflammatory immune interactions mediated by the equine Fc domain. More recently, the SARS-CoV-2 pandemic has generated renewed interest in prophylactic ruminant antibodies that potentially limit household viral transmission with at least one clinical trial currently underway (Uusküla et al., 2022). High antibody titers are key to the commercial viability of polyclonal preparations from ruminants, especially since the concentration of total IgGs in bovine colostrum drops rapidly from ∼50 g/L to ∼0.5 g/L in milk (Elfstrand et al., 2002) and antigen-specific antibodies may be less than 2% of total colostral antibodies following bovine vaccination (Chambers et al., 2022).

To improve antibody titers upon immunization, we selected IgG2a Fc domains for antigen fusion for two key reasons; first, this class of antibodies is poorly transported from serum into colostrum/milk and thus reduces the chance of undesired antigen contamination of downstream products (Nash et al., 1990). Second, in ruminant species the half-life of IgG2a in serum is frequently longer than for IgG1, which enables extended antigen persistence (Cervenak & Kacskovics, 2009). To our knowledge, this is the first demonstration of recombinant Fc fusion technology being used for the immunization of ruminant animals. However, the use of Fc fusion technology in other species has been observed to enhance immunization responses via well understood mechanisms. Fc domain engagement with FcRn receptors on host cells mediates the recycling of fusion proteins thus reducing the rate of catabolism and increasing antigen persistence (Rath et al., 2015). Our observed increases in serum titers correlates with those seen in studies with other species. For example, the 2.4-fold increase in serum titer at week 6 (three weeks after the first boost) is similar to the 1.6-fold increase in serum titer observed for RBD Fc-fused antigens in boosted mice (two weeks after the first boost) (Liu et al., 2020). An increase in both serum and neutralizing titer was also observed for IgG2a Fc fused to the full-length spike (but was not significant at p 0.05) but these increases were not as substantial as for RBD IgG2a Fc fusions. These differences may in part be due to the use of RBD antigens for serum titer analyses, which present only a proportion of the epitopes present within the full-length spike construct.

Further improvements to the immunogenicity of Fc-fused antigens might be made via multimerization, which enables the binding of the Fc domain to proinflammatory Fc receptors to promote antigen uptake and processing by antigen-presenting cells that initiate antibody responses (Nimmerjahn & Ravetch, 2008). Cui et al., recently demonstrated this approach by creating tetrameric Fc fusions with gp350 antigens from Epstein-Barr virus that were up to 25-fold more immunogenic in mice (Cui et al., 2013). We also expect manipulation of antibody responses (both magnitude and isotype distribution) could be achieved by investigating alternative antibody isotypes as Fc fusions, particularly since IgG1 fusions may traffic to the mammary gland to induce localized antibody responses in colostrum/milk. Deeper exploration of isotype bias will require the isolation of antigen specific antibodies, as we did not observe significant differences in isotype distribution in bulk colostrum or milk.

We also selected a second fusion, GM-CSF, as an immunofusion partner in an attempt to stimulate higher levels of antibodies following immunization. While some individual animals responded strongly, there was not a significant enhancement in neutralizing titer with the group sizes used in this study. GM-CSF has a broad range of immunological functions with known roles in the development of both antibody and cellular immunity, driven in part by the enhancement of antigen presenting cell function (Ushach & Zlotnik, 2016). Our findings suggest GM-CSF fusions should be revisited in future trials with greater statistical power. In either case, we note high animal-to-animal variation in response to immunization, particularly in the IgG2a and GM-CSF cohorts; a phenomenon well-established in sheep and other ruminant species (Hernández et al., 2003; Jouneau et al., 2020). In part, this is a consequence of the use of an outbred animal population in the study but could be resolved by using early serum titer testing to identify low-responding animals that could be subsequently removed from a dairy production herd. It is also likely that there is a genetic component to antibody immune responsiveness that may be exploitable in obtaining higher yielding production systems through selective breeding (McRae et al., 2015).

The rapid and continual evolution of SARS-CoV-2 variants has proved a challenge for the development of monoclonal antibody therapeutics. Polyclonal antibody preparations against SARS-CoV-2 may provide broader cross reactivity and we observe the neutralizing response generated in ovine colostrum covers several SARS-CoV-2 variants, albeit with slightly reduced efficacy against the omicron sublineage. These findings parallel observations in humans boosted with SARS-CoV-2 vaccines based on the spike protein antigen (Andrews et al., 2022; Lopez Bernal et al., 2021). However, further characterization of the colostral anti-SARS-CoV-2 antibody response is required. For instance, we have not tested neutralization potency with live virus. There also remain outstanding questions as to the pharmacokinetics of the injected antigens. While the IgG1 subclass is preferentially trafficked into colostrum/milk, we have not ruled out the presence of IgG2a Fc fusions in the colostrum though concentrations are likely very low, if present at all. Tackling this question will require more sensitive methods than the ELISA approaches developed here, potentially using mass spectrometry to screen for RBD peptide presence in colostrum/milk. Nonetheless, the use of fusion proteins to stimulate antibody responses in ruminants may enable new pathways for efficient hyperimmune milk generation as well as in the development of next generation vaccines against ruminant pathogens.

## Conclusion

Technologies to increase the levels of circulating antibodies in ruminant animals are important for biotechnology and animal health applications. We explored the potency of immunomodulatory fusion proteins IgG2a Fc and GM-CSF to enhance the antibody response upon immunization. Our results show that fusion of ovine IgG2a Fc domains to the SARS-CoV-2 RBD model antigen elicits significantly higher neutralizing antibody titers in sheep than with the RBD antigen alone. Fusion of GM-CSF cytokines to the RBD was unsuccessful at increasing neutralizing titers, despite the observation of several high-responding individual animals. We further demonstrate that antibodies resulting from the fusion proteins are trafficked into milk following lambing and due to the polyclonal nature of the response are able to neutralize several SARS-CoV-2 variants. We suggest that the action of our antigenic fusions is mediated by Fc receptor engagement and that future designs could further improve immunogenicity via multimerization or the use of alternative isotypes as Fc domain fusions.

## Acknowledgements

We would like to thank the staff of the animal facility (AgResearch, Hamilton, New Zealand) for their help with trial management. We also thank Sophie Cook for her assistance with the development of reagents used for ovine serum titering and Harold Henderson for his assistance with statistical analyses. The RuaTech R&D contribution was co funded under Callaghan Project Grant RTECD 2003 and R&D Loan RTECD 2001. WK was supported by a Royal Society Te Apārangi Marsden Fund Fast-Start grant (19-FRI-002).

## Author contributions

WK, AW, RC, GS, AH, MF and SH conceived and designed the research; GJ, KK, OW, GS, JP and WK performed the research; GJ, KK and WK wrote the first draft of the article, and all authors discussed and contributed to the manuscript.

