## Additional File 1; Supplementary information for "Immunogenic fusion proteins induce neutralizing SARS-CoV-2 antibodies in the serum and milk of sheep"

### Supplementary Tables

**Table S1.** Sequences for SARS-CoV-2 antigen fusions

| | |
|---|---|
| >RBD | METGLRWLLLVAVLKGVQCRVQPTESIVRFPNITNLCPFGEVFNATRFASVYAWNR<br>KRISNCVADYSVLYNSASFSTFKCYGVSPTKLNDLCFTNVYADSFVIRGDEVRQIAPG<br>QTGKIADYNYKLPDDFTGCVIAWNSNNLDSKVGGNYNYLYRLFRKSNLKPFERDIST<br>EIYQAGSTPCNGVEGFNCYFPLQSYGFQPTNGVGYQPYRVVVLSFELLHAPATVCGP<br>KKSTNLVKNKCVNFNFNGGGGGSHHHHHH |
| >RBD IgG2a Fc | METGLRWLLLVAVLKGVQCRVQPTESIVRFPNITNLCPFGEVFNATRFASVYAWNR<br>KRISNCVADYSVLYNSASFSTFKCYGVSPTKLNDLCFTNVYADSFVIRGDEVRQIAPG<br>QTGKIADYNYKLPDDFTGCVIAWNSNNLDSKVGGNYNYLYRLFRKSNLKPFERDIST<br>EIYQAGSTPCNGVEGFNCYFPLQSYGFQPTNGVGYQPYRVVVLSFELLHAPATVCGP<br>KKSTNLVKNKCVNFNFNGDYSKCSKPPCVSRPSVFIFPPKPKDSLMITGTPEVTCVVV<br>DVGQGDPEVQFSWFVDNVEVRTARTKPREEQFNSTFRVVSALPIQHDHWTGGKEFK<br>CKVHSKGLPAPIVRTISRAKGQAREPQVYVLAPPQEELSKSTLSVTCLVTGFYPDYIA<br>VEWQRARQPESEDKYGTTTSQLDADGSYFLYSRLRVDKSSWQRGDTYACVVMHEA<br>LHNHYTQKSISKPPGKGGGGSHHHHHH |
| >RBD GM-CSF | METGLRWLLLVAVLKGVQCRVQPTESIVRFPNITNLCPFGEVFNATRFASVYAWNR<br>KRISNCVADYSVLYNSASFSTFKCYGVSPTKLNDLCFTNVYADSFVIRGDEVRQIAPG<br>QTGKIADYNYKLPDDFTGCVIAWNSNNLDSKVGGNYNYLYRLFRKSNLKPFERDIST<br>EIYQAGSTPCNGVEGFNCYFPLQSYGFQPTNGVGYQPYRVVVLSFELLHAPATVCGP<br>KKSTNLVKNKCVNFNFNGGGGGSAPTRQPSPVTRPWQHVDAIKEALSLLNDSTDTA<br>AVMDETVEVVSEMFDSQEPTCLQTRLELYKQGLRGSLTSLTGSLTMMASHYKKHCP<br>PTQETSCETQIITFKSFKENLKDFLFIIPFDCWEPVQKGGGGSHHHHHH |
| >Spike IgG2a Fc | METGLRWLLLVAVLKGVQCFVFLVLLPLVSSQCVNLTTRTQLPPAYTNSFTRGVYY<br>PDKVFRSSVLHSTQDLFLPFFSNVTWFHAIHVSGTNGTKRFDNPVLPFNDGVYFASTE<br>KSNIIRGWIFGTTLDSKTQSLLIVNNATNVVIKVCEFQFCNDPFLGVYYHKNNKSWM<br>ESEFRVYSSANNCTFEYVSQPFLMDLEGKQGNFKNLREFVFKNIDGYFKIYSKHTPIN<br>LVRDLPQGFSALEPLVDLPIGINITRFQTLLALHRSYLTPGDSSSGWTAGAAAYYVGY<br>LQPRTFLLKYNENGTITDAVDCALDPLSETKCTLKSFTVEKGIYQTSNFRVQPTESIVR<br>FPNITNLCPFGEVFNATRFASVYAWNRKRISNCVADYSVLYNSASFSTFKCYGVSPT<br>KLNDLCFTNVYADSFVIRGDEVRQIAPGQTGKIADYNYKLPDDFTGCVIAWNSNNL<br>DSKVGGNYNYLYRLFRKSNLKPFERDISTEIYQAGSTPCNGVEGFNCYFPLQSYGFQP<br>TNGVGYQPYRVVVLSFELLHAPATVCGPKKSTNLVKNKCVNFNFNGLTGTGVLTES<br>NKKFLPFQQFGRDIADTTDAVRDPQTLEILDITPCSFGGVSVITPGTNTSNQVAVLYQ<br>GVNCTEVPVAIHADQLTPTWRVYSTGSNVFQTRAGCLIGAEHVNNSYECDIPIGAGI<br>CASYQTQTILRDYSKCSKPPCVSRPSVFIFPPKPKDSLMITGTPEVTCVVVDVGQGDP<br>EVQFSWFVDNVEVRTARTKPREEQFNSTFRVVSALPIQHDHWTGGKEFKCKVHSKG<br>LPAPIVRTISRAKGQAREPQVYVLAPPQEELSKSTLSVTCLVTGFYPDYIAVEWQRAR<br>QPESEDKYGTTTSQLDADGSYFLYSRLRVDKSSWQRGDTYACVVMHEALHNHYTQ<br>KSISKPPGKGGGGSHHHHHH |

**Table S2.** Power law coefficients, Pearson correlation values and associated p values for serum titers compared to serum neutralizing titers for each animal cohort

| | Week 6 | | | Week 9 | | |
|---|---|---|---|---|---|---|
| **Treatment** | Slope Coefficient[#] | Correlation r | p value | Slope Coefficient[#] | Correlation r | p value |
| **RBD** | 0.454 (0.172-0.735) | 0.80 | 0.031 | 0.498 (0.126-0.869) | 0.77 | 0.044 |
| **RBD IgG2 Fc** | 1.266 (0.619-1.914) | 0.86 | 0.013 | 1.062 (0.468-1.656) | 0.79 | 0.034 |
| **RBD GM CSF** | 1.166 (0.716-1.617) | 0.98 | 0.0002 | 0.972 (0.542-1.402) | 0.99 | <0.0001 |
| **Spike Low** | 0.714 (0.289-1.138) | 0.95 | 0.0012 | 0.759 (0.120-1.397) | 0.89 | 0.0076 |
| **Spike High** | 0.839 (0.454-1.225) | 0.88 | 0.0083 | 0.882 (0.515-1.249) | 0.87 | 0.012 |
| **Spike IgG2 Fc** | 0.556 (0.155-0.957) | 0.67 | 0.1 | 0.464 (-0.004-0.933) | 0.61 | 0.14 |

[#]Brackets indicate 95% confidence limits from power law fitting

**Table S3.** Statistical testing of serum viral neutralization. Means were obtained for viral neutralization titers and adjusted for mean serum titre for each treatment.

| | **Week 6** | | **Week 9** | |
|---|---|---|---|---|
| **Treatment** | Mean | Group[#] | Mean | Group[#] |
| **RBD** | 1.86 | CD | 1.66 | B |
| **RBD IgG2 Fc** | 1.88 | CD | 1.62 | AB |
| **RBD GM CSF** | 2.09 | D | 1.87 | B |
| **Spike Low** | 1.42 | A | 1.35 | A |
| **Spike High** | 1.46 | AB | 1.39 | A |
| **Spike IgG2 Fc** | 1.75 | BC | 1.67 | B |

[#]Means with the same letter (within week) are not significantly different at $p \leq 0.05$

**Table S4.** Power law coefficients, correlation values and associated p values for serum titers compared to milk titers for days 0, 4, 16, and 32 after lambing.

| Time | Slope Coefficient[#] | Correlation r | p value |
|---|---|---|---|
| **Day 0** | 0.306 (0.036-0.577) | 0.54 | 0.029 |
| **Day 4** | 0.543 (0.155-0.931) | 0.63 | 0.0095 |
| **Day 16** | 0.897 (0.642-1.151) | 0.90 | <0.0001 |
| **Day 32** | 0.647 (0.361-0.934) | 0.80 | 0.0003 |

[#]Brackets indicate 95% confidence limits from power law fitting

### Supplementary Figures

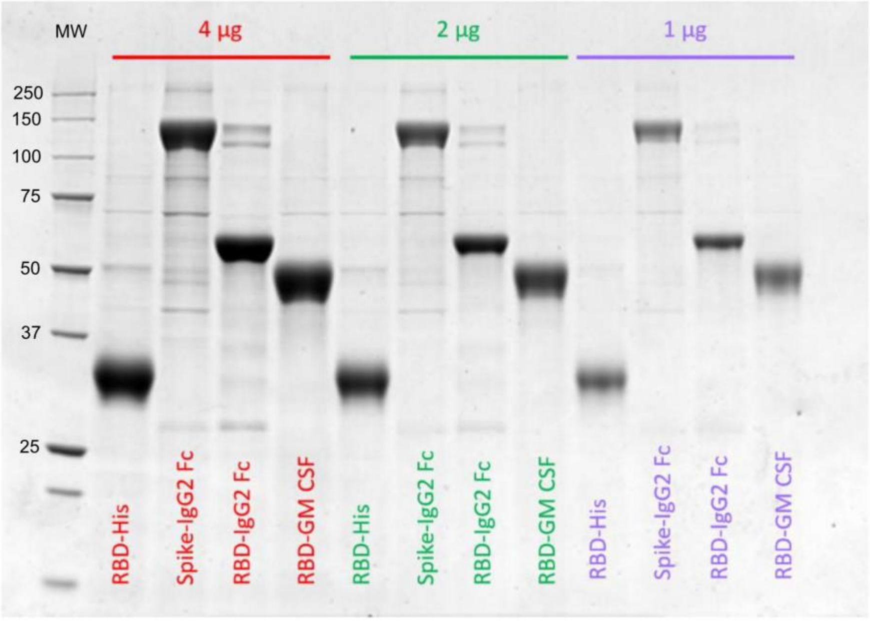


**Figure S1.** SDS-PAGE gel showing purified vaccine antigens. Protein masses loaded are indicated on the gel. MW = Precision Plus molecular weight ladder (Biorad, USA).

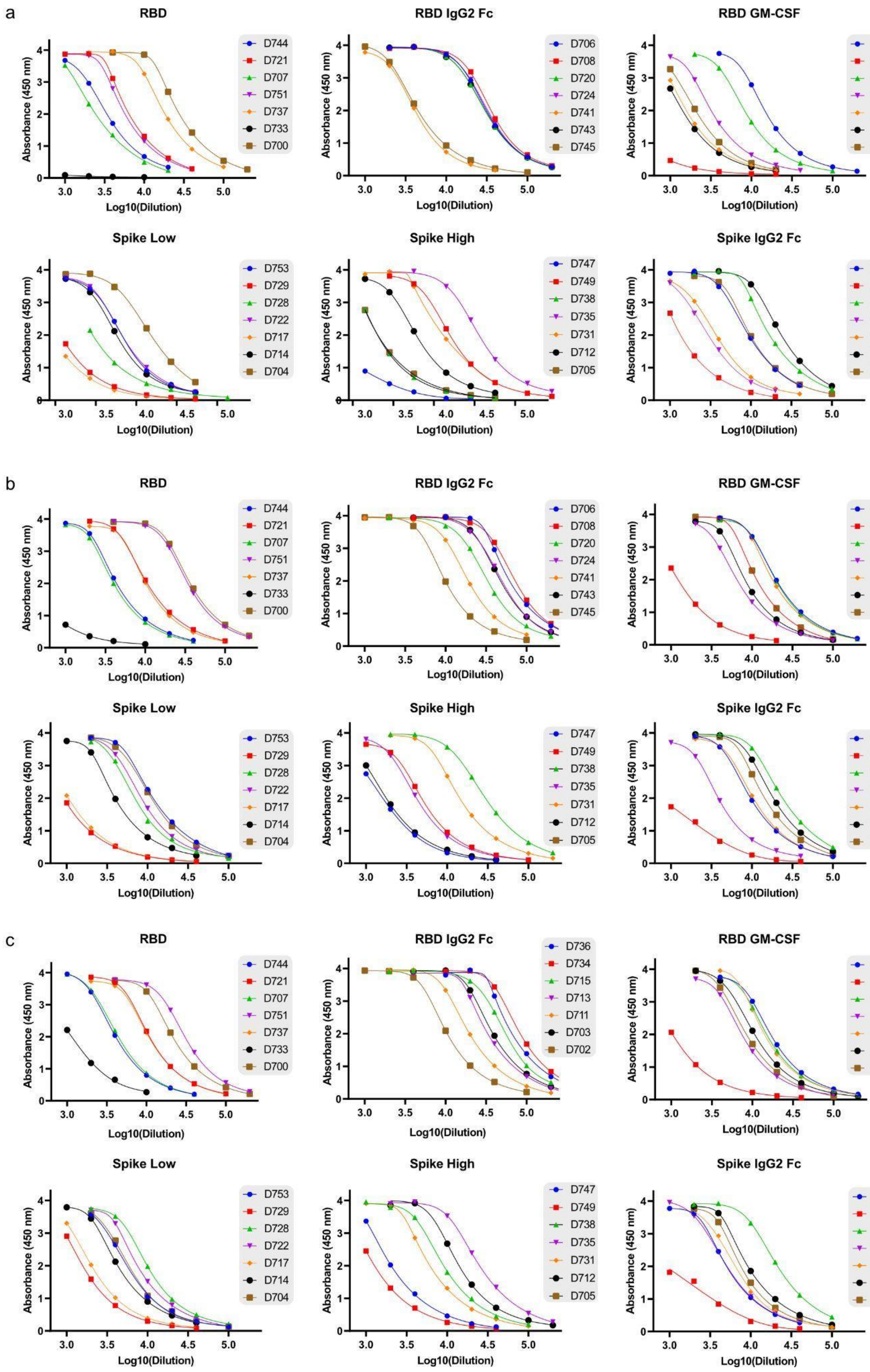
a
RBD
D744
D721
D707
D751
D737
D733
D700
RBD IgG2 Fc
D706
D708
D720
D724
D741
D743
D745
RBD GM-CSF
D755
D754
D727
D725
D718
D716
D701
Spike Low
D753
D729
D728
D722
D717
D714
D704
Spike High
D747
D749
D738
D735
D731
D712
D705
Spike IgG2 Fc
D736
D734
D715
D713
D711
D703
D702
Absorbance (450 nm)
Log10(Dilution)
b
RBD
D744
D721
D707
D751
D737
D733
D700
RBD IgG2 Fc
D706
D708
D720
D724
D741
D743
D745
RBD GM-CSF
D755
D754
D727
D725
D718
D716
D701
Spike Low
D753
D729
D728
D722
D717
D714
D704
Spike High
D747
D749
D738
D735
D731
D712
D705
Spike IgG2 Fc
D736
D734
D715
D713
D711
D703
D702
Absorbance (450 nm)
Log10(Dilution)
c
RBD
D744
D721
D707
D751
D737
D733
D700
RBD IgG2 Fc
D736
D734
D715
D713
D711
D703
D702
RBD GM-CSF
D755
D754
D727
D725
D718
D716
D701
Spike Low
D753
D729
D728
D722
D717
D714
D704
Spike High
D747
D749
D738
D735
D731
D712
D705
Spike IgG2 Fc
D706
D708
D720
D724
D741
D743
D745
Absorbance (450 nm)
Log10(Dilution)

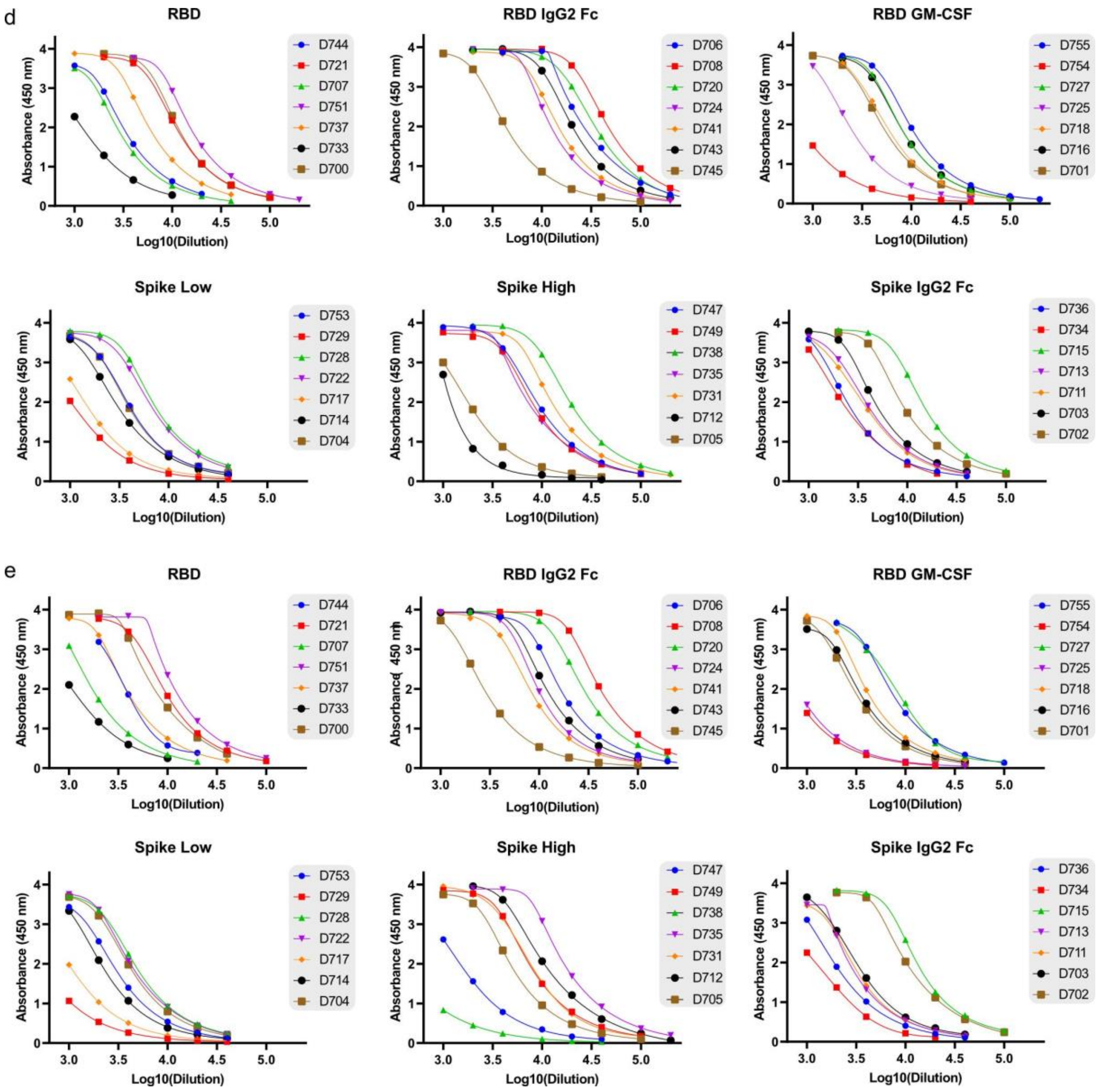


**Figure S2.** Total antibody titers for sera samples from weeks 3 (a), 6 (b), 9 (c), 12 (d) and 16 (e). Measurements were made in duplicate by determining serum antibody binding to Wuhan RBD protein and the resulting curves were fitted with sigmoidal functions to obtain IC50 values.

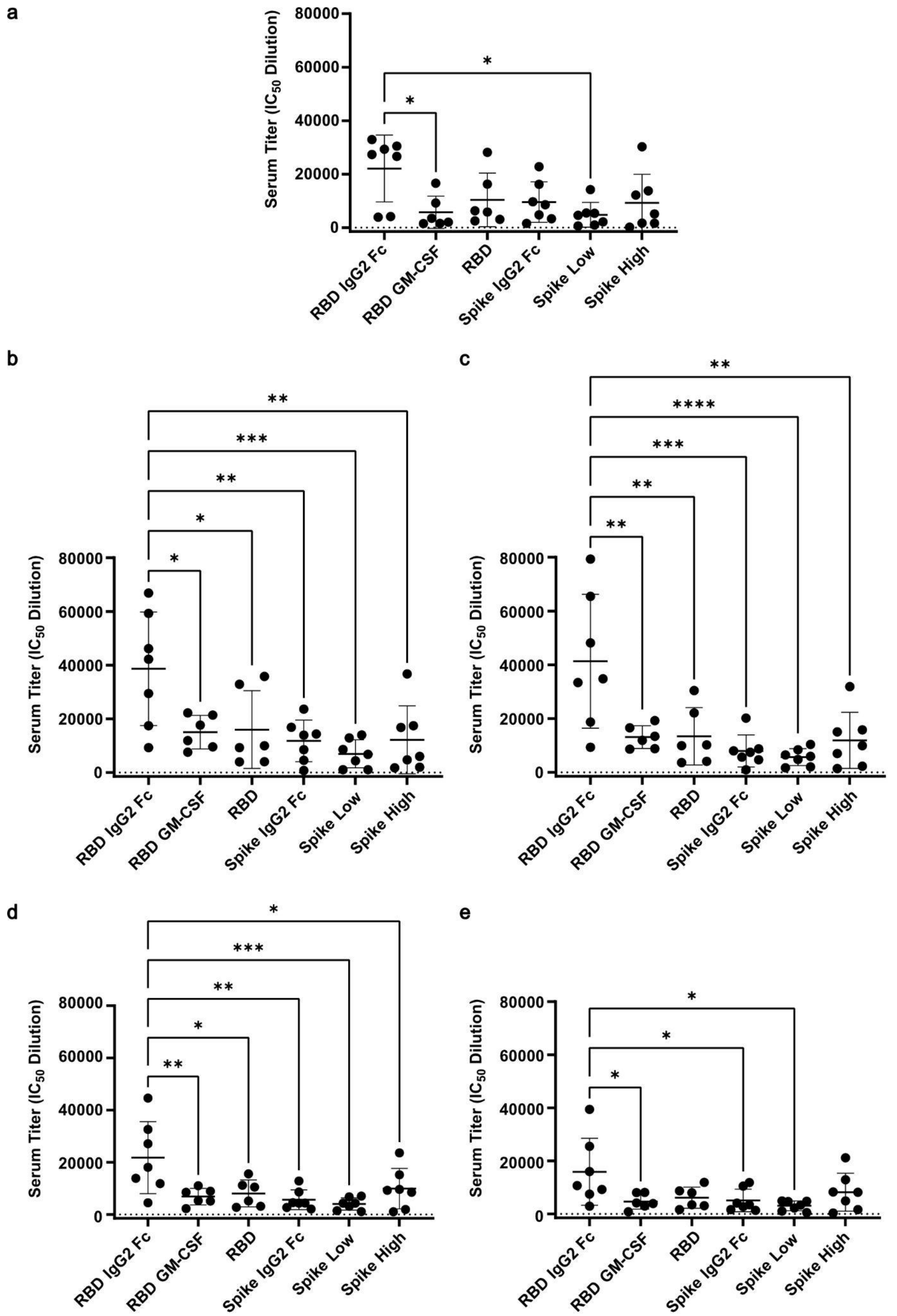


**Figure S3.** IC50 plots for serum titers at week a) 3; b) 6; c) 9; d) 12; e) 16. p-values were determined by ANOVA (*≤0.05, **≤0.01, **≤0.001, and ***<0.0001). Error bars represent the standard error for each group.

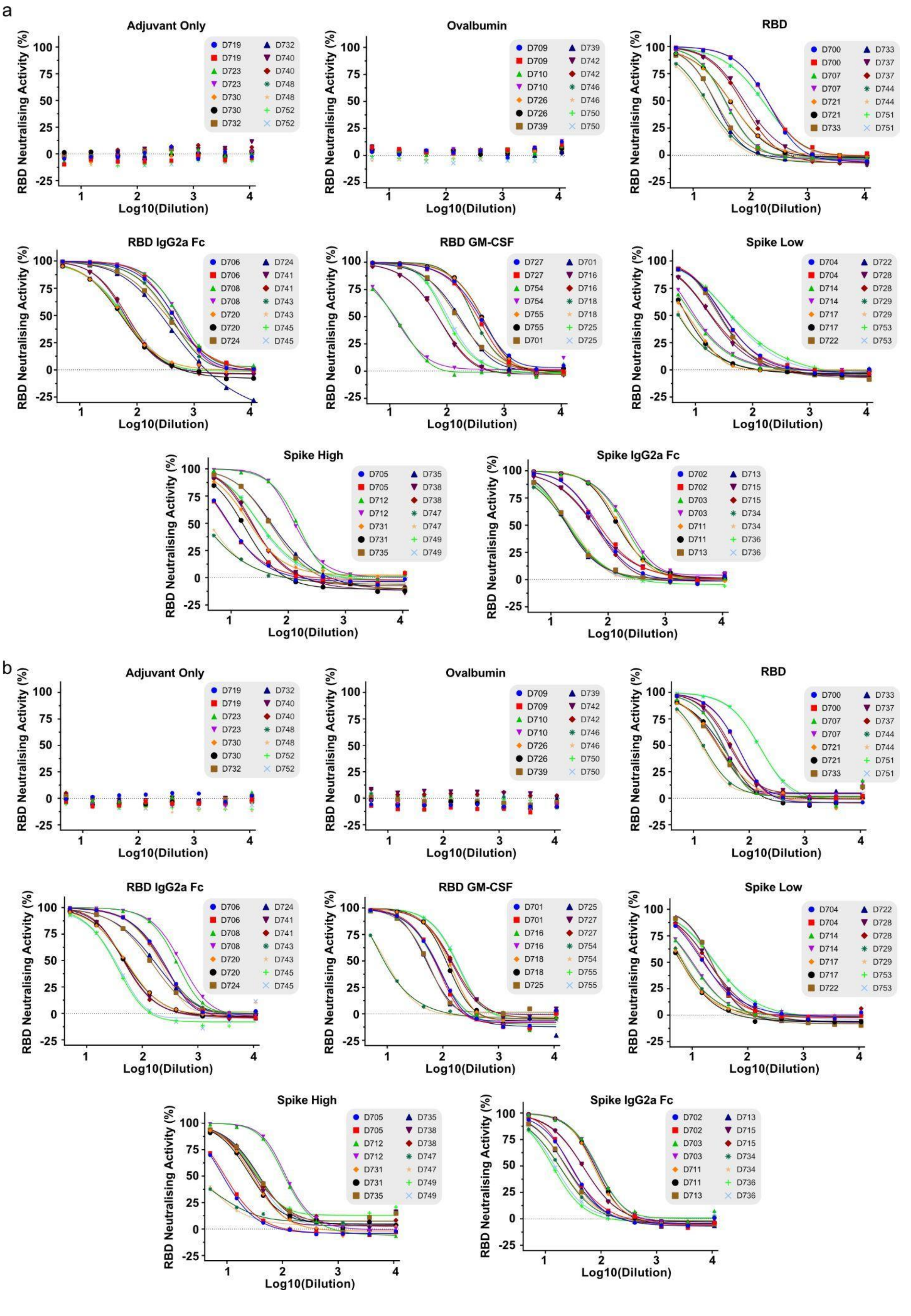
a
Adjuvant Only
RBD Neutralising Activity (%)
Log10(Dilution)
D719 D719 D723 D723 D730 D730 D732 D732 D740 D740 D748 D748 D752 D752
Ovalbumin
D709 D709 D710 D710 D726 D726 D739 D739 D742 D742 D746 D746 D750 D750
RBD
D700 D700 D707 D707 D721 D721 D733 D733 D737 D737 D744 D744 D751 D751
RBD IgG2a Fc
D706 D706 D708 D708 D720 D720 D724 D724 D741 D741 D743 D743 D745 D745
RBD GM-CSF
D727 D727 D754 D754 D755 D755 D701 D701 D716 D716 D718 D718 D725 D725
Spike Low
D704 D704 D714 D714 D717 D717 D722 D722 D728 D728 D729 D729 D753 D753
Spike High
D705 D705 D712 D712 D731 D731 D735 D735 D738 D738 D747 D747 D749 D749
Spike IgG2a Fc
D702 D702 D703 D703 D711 D711 D713 D713 D715 D715 D734 D734 D736 D736
b
Adjuvant Only
D719 D719 D723 D723 D730 D730 D732 D732 D740 D740 D748 D748 D752 D752
Ovalbumin
D709 D709 D710 D710 D726 D726 D739 D739 D742 D742 D746 D746 D750 D750
RBD
D700 D700 D707 D707 D721 D721 D733 D733 D737 D737 D744 D744 D751 D751
RBD IgG2a Fc
D706 D706 D708 D708 D720 D720 D724 D724 D741 D741 D743 D743 D745 D745
RBD GM-CSF
D701 D701 D716 D716 D718 D718 D725 D725 D727 D727 D754 D754 D755 D755
Spike Low
D704 D704 D714 D714 D717 D717 D722 D722 D728 D728 D729 D729 D753 D753
Spike High
D705 D705 D712 D712 D731 D731 D735 D735 D738 D738 D747 D747 D749 D749
Spike IgG2a Fc
D702 D702 D703 D703 D711 D711 D713 D713 D715 D715 D734 D734 D736 D736

**Figure S4.** Neutralizing activity within sera from individual animals at a) 6 and b) 9 weeks post-vaccination as measured by sVNT assay. Measurements were made in duplicate and indicate the neutralizing ability of sera against the Wuhan RBD protein. All curves were fitted with sigmoidal functions to obtain IC50 values.

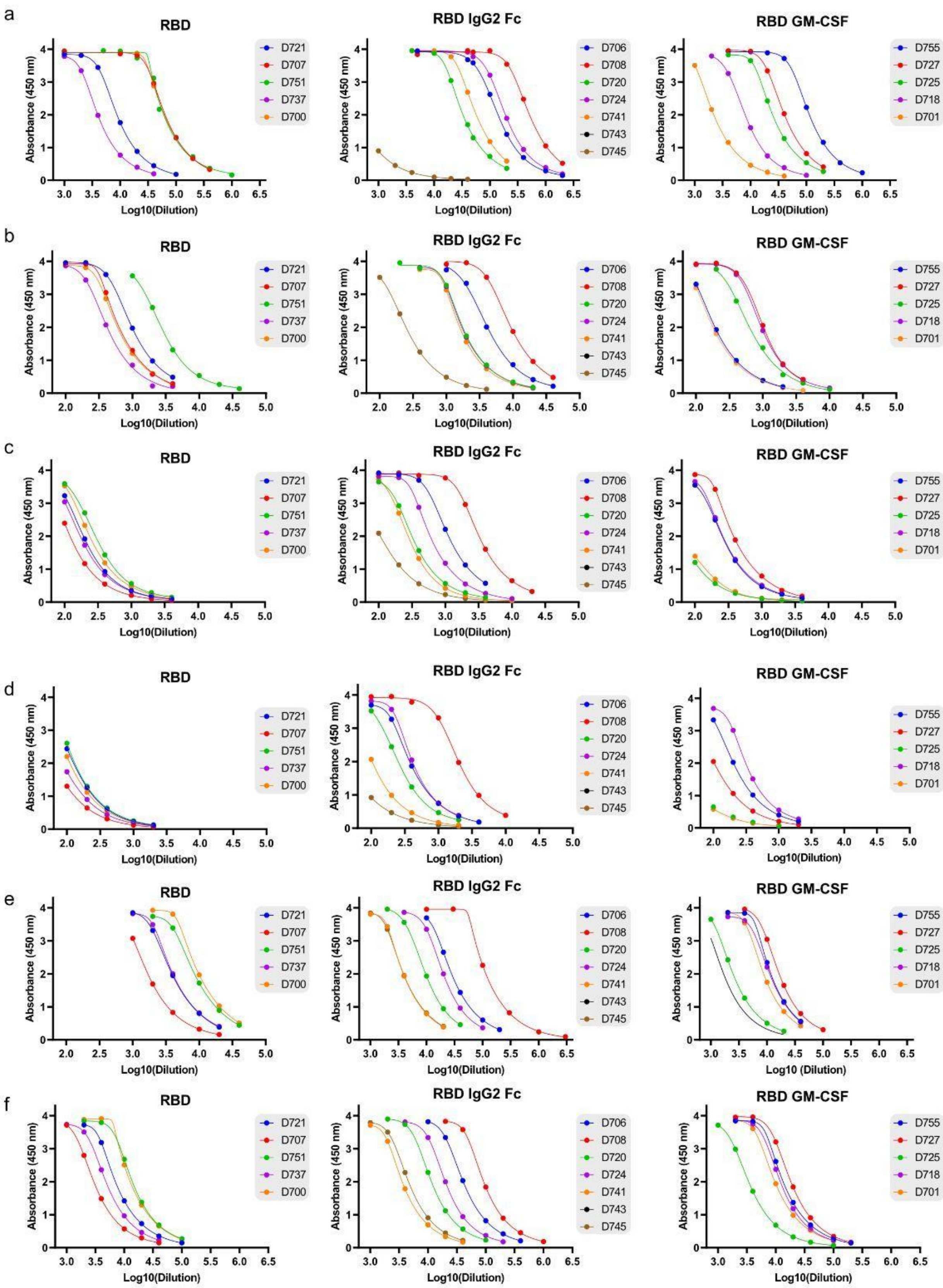
a
b
c
d
e
f
RBD
RBD IgG2 Fc
RBD GM-CSF
Absorbance (450 nm)
Log10(Dilution)
Log10 (Dilution)
D721
D707
D751
D737
D700
D706
D708
D720
D724
D741
D743
D745
D755
D727
D725
D718
D701

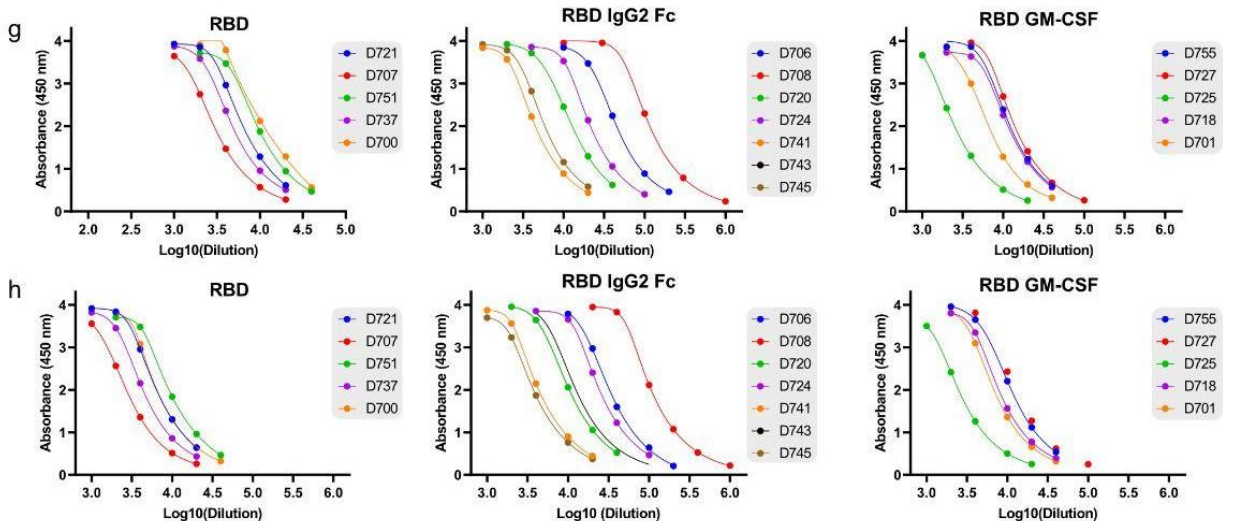


**Figure S5.** Total antibody titers from the milk trial extension measured in milk at day 0 (a), 4 (b), 16 (c), and 32 (d) and serum at also at days 0 (e), 4 (f), 16 (g), and 32 (h). Measurements were made in duplicate by determining antibody binding to Wuhan RBD protein and the resulting curves were fitted with sigmoidal functions to obtain IC50 values.

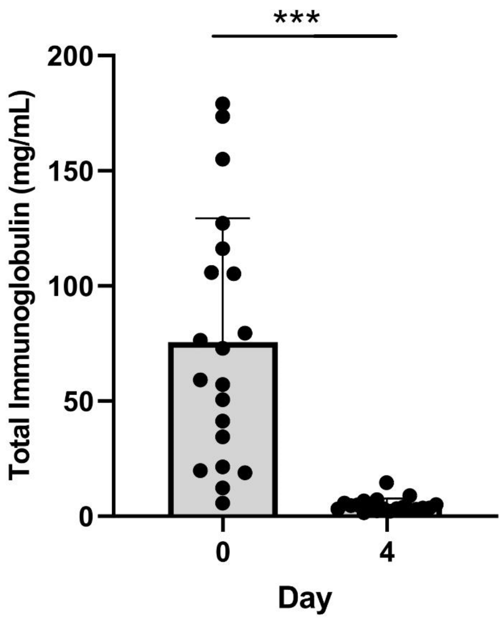


**Figure S6.** Total antibody concentration in colostrum at days 0 and 4 as measured by mass spectrometry. Error bars show the standard error for each group. Student's T Test *** p<0.0001

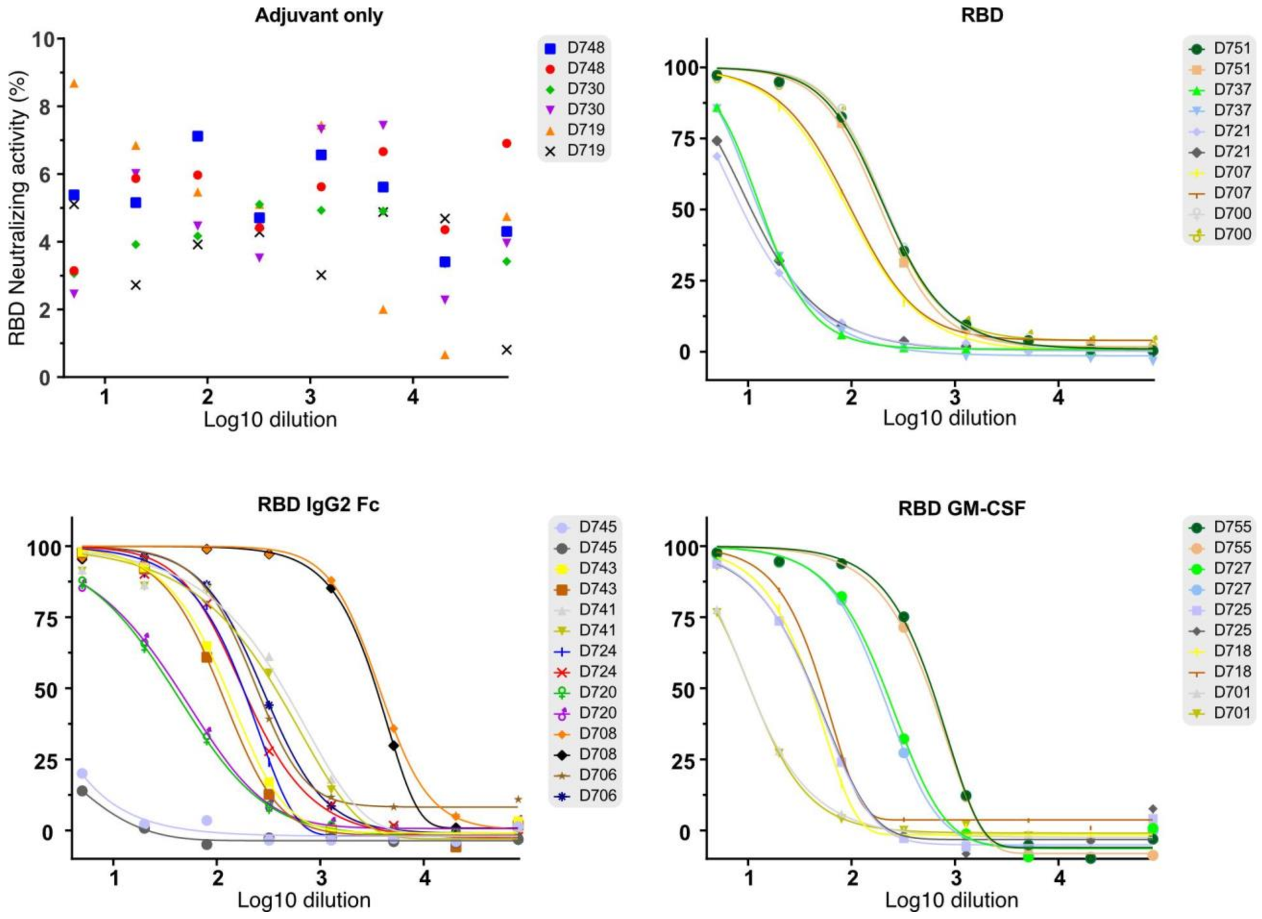

**Figure S7.** Neutralizing activity within colostrum from individual animals at day 0 post-lambing as measured by sVNT assay. Measurements were made in duplicate and indicate the neutralizing ability of sera against the Wuhan RBD protein. All curves were fitted with sigmoidal functions to obtain IC50 values.
